# Live-cell imaging reveals nutrient-dependent dynamics of ER-mitochondria contact formation via PDZD8

**DOI:** 10.1101/2025.04.17.649323

**Authors:** Saeko Aoyama-Ishiwatari, Koki Nakamura, Takahiro Nagao, Yusuke Hirabayashi

## Abstract

Cells adapt to changes in nutrient conditions by reorganizing organelle activities. Mitochondria-endoplasmic reticulum contact sites (MERCS) have been proposed to play an essential role in this adaptive response. However, the lack of suitable tools for detecting these contact sites in living cells has hindered the studies of the mechanisms regulating MERCS in response to nutritional changes. Here, we establish a novel cell line, MERCdRED, which expresses a ddFP-based fluorescent MERCS sensor. Using correlative light-electron microscopy, we show that the MERCdRED signal accurately labels the contact sites as defined by electron microscopy. Live imaging of MERCdRED cells revealed that large MERCS are more stable than smaller ones. Furthermore, in combination with knockout of the ER-mitochondria tethering protein PDZD8, we revealed that nutrient deprivation reduces MERCS area in a PDZD8-dependent manner. This new tool, together with our findings, will contribute to a better understanding of the molecular basis of cellular metabolism.

**Summary Statement:** We established a novel cell line enabling live imaging of MERCS, which revealed their dynamic nature and demonstrated nutrient deprivation-induced MERCS reduction along with its underlying mechanisms.

## Introduction

Ultrastructural analysis using electron microscopy (EM) has revealed that organelles form contact sites where their membranes are juxtaposed within tens of nanometers. Mitochondria-endoplasmic reticulum contact sites (MERCS) are the most abundant inter-organelle contacts in many cell types and serve as hubs for various cellular biochemical reactions (Csordás et al., 2006; Prinz et al., 2020; Wu et al., 2018). The formation of MERCS requires protein complexes that tether the ER and mitochondria. The identification of multiple tethering complexes suggests that distinct protein complexes play specialized roles in regulating MERCS (Aoyama-Ishiwatari & Hirabayashi, 2021).

Changes in MERCS formation in response to nutrient excess have been investigated as a potential mechanism underlying ER and mitochondrial dysfunction in obese animal models. In these studies, MERCS abundance has been assessed using widely employed methods such as EM observations of fixed tissues and biochemical approaches for isolating contact sites (Arruda et al., 2014; Parlakgül et al., 2024; Theurey et al., 2016; Tubbs et al., 2014). Nevertheless, contradictory findings have been reported, and how MERCS respond to nutritional changes in living cells remains unclear.

While fluorescence microscopy has been applied to image MERCS in living cells, its limited resolution has hampered precise detection of the contact site. To address this limitation, several studies have reported genetically encoded fluorescent probes for specific labeling of MERCS. Among these, the reversibility of low-affinity dimerization-dependent detection systems, such as the Förster resonance energy transfer (FRET) (Csordás et al., 2010) and the dimerization-dependent fluorescent protein (ddFP) (Abrisch et al., 2020; Naon et al., 2016; Naón et al., 2023; Sakai et al., 2021), has made them promising tools. However, previous studies using these systems have relied on transient gene expression methods, which result in significant variations in expression levels of these probes among individual cells. As a result, quantitative analysis of changes in MERCS remains challenging.

In this study, we developed a novel MERCS detection system by establishing a clonal cell line stably expressing ddFP, enabling consistent and quantitative evaluation of MERCS. Using this cell line, we revealed that the ER protein PDZD8, an essential tether of MERCS (Hirabayashi et al., 2017; Nakamura et al., 2025), facilitates MERCS formation under nutrient-rich conditions. This new tool and our findings provide insights for elucidating the molecular mechanisms underlying the regulation of MERCS dynamics.

## Results & Discussion

### Single cell cloning of a stable cell line expressing MERCdRED system

To investigate dynamic changes of MERCS amount in response to nutritional changes in living cells, we utilized the ddFP system consisting of quenched or non-fluorescent proteins that emit fluorescence upon dimerization (Alford et al., 2012; Ding et al., 2015). To adapt this system for MERCS visualization in live cells, we fused the quenched red fluorescent component (RA) to human Sec61β and the nonfluorescent component (GB) to human Tomm20 to anchor them to the cytosolic sides of the ER membrane (ERM) and the outer mitochondrial membrane (OMM), respectively (**Fig. 1A, B**). With the aim of elucidating the role of the ER-mitochondria tethering protein PDZD8 in MERCS regulation upon nutritional changes, a plasmid encoding the RA part and the GB part connected with a sequence for the self-cleaving peptide P2A (Tomm20-GB-P2A-RA-Sec61β) was introduced into *Pdzd8*^f/f^::Cre^ERT2^ mouse embryonic fibroblasts (MEFs), which allows tamoxifen-inducible conditional knockout of PDZD8 (Nakamura et al., 2025). Given that MERCS detection by ddFP relies on affinity-dependent interactions, variations in the expression levels of ddFP component proteins could significantly impact fluorescence intensity variance, regardless of the actual MERCS area. Thus, we established a stable cell line with uniform expression of the proteins for quantitative analysis and termed it “MERCdRED cell”. The Tomm20-GB-P2A-RA-Sec61β-encoding sequence was integrated into the genome via lentiviral infection, followed by single-cell cloning (**Fig. 1C**). Among proliferative clones, the one exhibiting the highest red fluorescence intensity was selected using confocal microscopy. While non-mitochondrial autofluorescence was observed in both control and MERCdRED-cells, strong red fluorescence was specifically detected on mitochondria in MERCdRED cells (**Fig. 1D, E**). Notably, the red fluorescence on mitochondria was observed in close proximity to the ER (**Fig. 1E**). Note that these red signals, referred to as MERCdRED fluorescence, were only visualized in live cells but not in fixed cells with paraformaldehyde and glutaraldehyde (**Fig. 1D**). Quantification of red fluorescence in mitochondrial regions revealed a significantly higher signal intensity in MERCdRED cells compared to control cells (**Fig. 1F**), indicating the successful dimerization of Tomm20-GB and GFP-Sec61β on mitochondria in the MERCdRED cell line.

**Figure 1.**
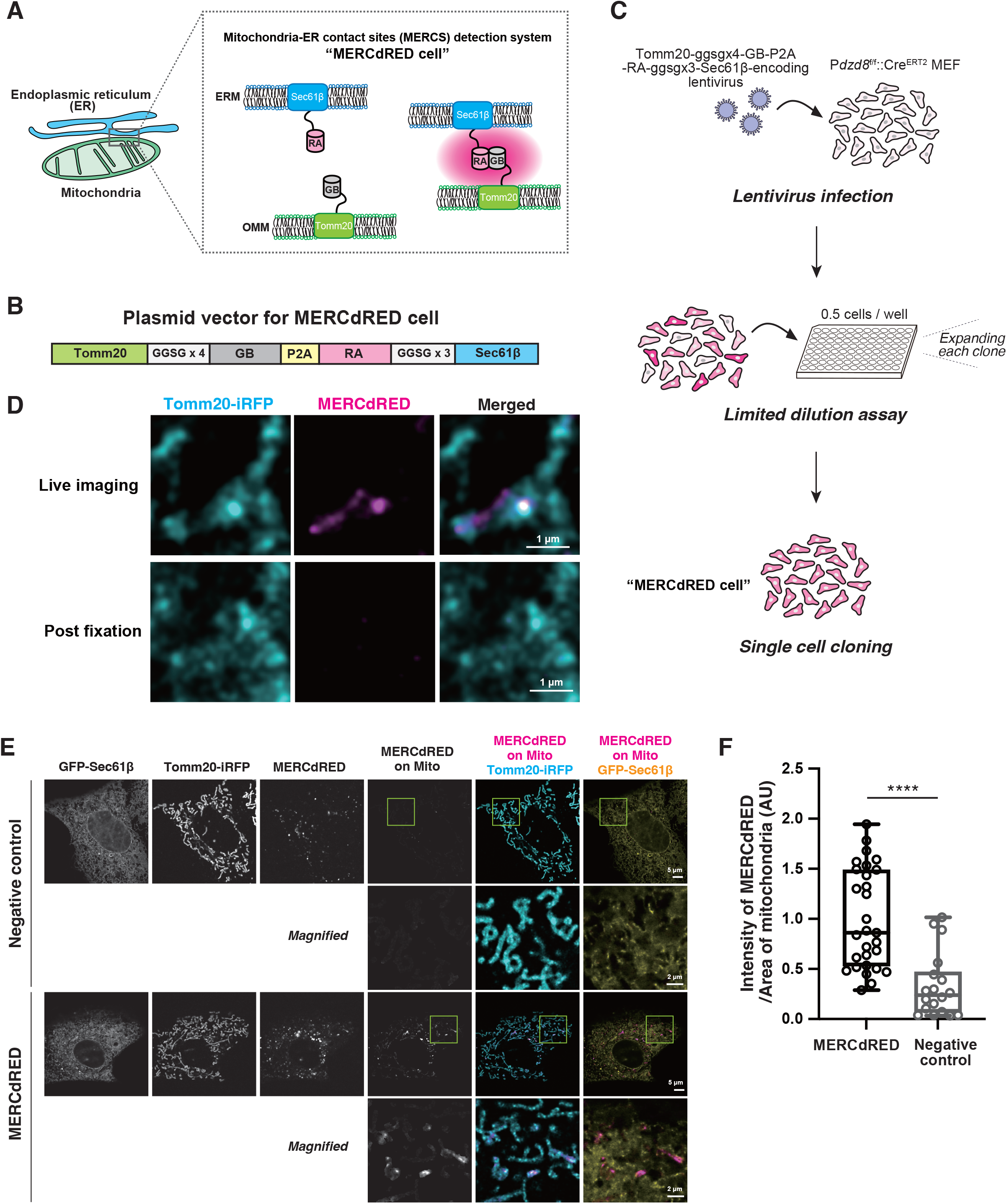
Establishment of the MERCdRED system for visualizing MERCS in live cells. **(A)** Schematic representation of the MERCdRED system. **(B)** Diagram of the MERCdRED-expressing vector. Two components of the MERCdRED system, Tomm20-GB and RA-Sec61β, were co-expressed from a single vector using a P2A sequence, which enables post-translational self-cleavage. **(C)** Diagram illustrating the establishment of the MERCdRED cell line. **(D)** Comparison of MERCdRED signals between live cells and cells fixed with 2% paraformaldehyde (PFA) and 0.5% glutaraldehyde (GA). **(E)** Representative live images of the MERCdRED cells (MERCdRED shown in magenta) transfected with plasmids encoding an ER marker GFP-Sec61β (yellow) or a mitochondrial marker Tomm20-iRFP (cyan). The boxed regions of the top panels are shown at higher magnification in the corresponding lower panels. **(F)** Quantification of the MERCdRED intensity on mitochondria in images obtained as described (E). The data are presented as individual points on box plots, with the center indicating the median, and the 25th and 75th percentiles represented by the box. Whiskers extend to the minimum and maximum values. n = 29, 18 cells for the MERCdRED cells and the control cells (negative control) from five independent experiments. Statistical analysis was performed using two-tailed Student’s t-test. *\*\*\*\*P* < 0.0001

### Accurate labeling of MERCS by MERCdRED fluorescence demonstrated by correlative light-electron microscopy

To determine whether MERCdRED signals on mitochondria reliably represent MERCS, we performed the three-dimensional (3D) correlative light-electron microscopy (CLEM) analysis in live cells (Live-3D CLEM) using the MERCdRED cell line (**Fig. 2A**). First, signals from MERCdRED fluorescence overlapped with the mitochondrial marker Tomm20-iRFP were detected by confocal microscopy in living cells. The cells were immediately fixed, stained, dehydrated, and embedded in a resin. Then serial 50 nm-thick sections of the cells were imaged by a field-emission scanning EM. Using mitochondria reconstructed into a 3D image from ten serial EM slices as landmarks, the areas imaged by confocal microscopy were re-identified in EM images (**Fig. 2B**). Notably, the MERCdRED signals were highly colocalized with 3D-reconstructed MERCS where the OMM and the ER membrane were within 15 nm of each other (**Fig. 2B, C**). Due to disproportionate shrinkage of the cell during the fixation and dehydration steps, peripheral areas of the confocal and EM images were not perfectly aligned. Therefore, we focused on regions where mitochondria imaged by the confocal microscopy and EM are well aligned for further analysis. In those regions, accuracy and precision of MERCdRED signals corresponding to MERCS were calculated from chunk-averaged images, which were generated by averaging the proximity-mapped images within each 250 nm square chunk (**Fig. 2D**). Both accuracy and precision values ranged largely from 0.7 to 0.9, which are several folds higher compared to those obtained when the MERCdRED chunk-averaged images were randomized (**Fig. 2E**). The apparent false-positive and false-negative signals of MERCdRED are likely attributable to the time gap between live-cell imaging and fixation, differences in optical section thickness between confocal imaging and 3D EM reconstruction, and nonspecific fluorescence of MERCdRED. Additionally, some small MERCS were not detectable, presumably due to the relatively weak fluorescence of MERCdRED. Overall, the MERCdRED system effectively visualized MERCS with a 15 nm intermembrane gap.

**Figure 2.**
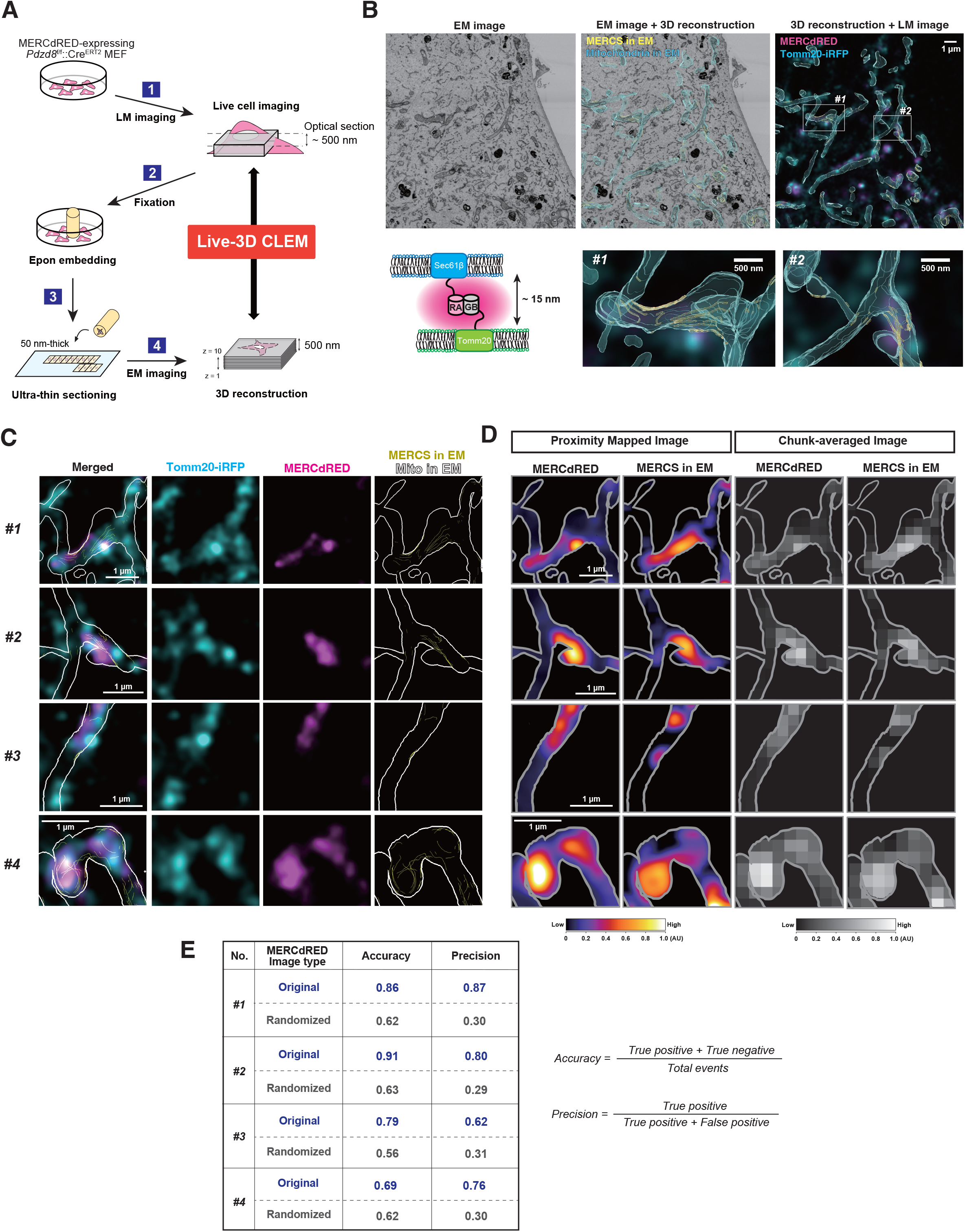
Correlative light-electron microscopy analysis demonstrates that MERCdRED signals correspond to MERCS identified in EM. **(A)** Schematic representation of Live-3D CLEM analysis. The MERCdRED cells were imaged live (shown as light microscopy (LM) imaging), and subsequently fixed with paraformaldehyde (PFA), glutaraldehyde (GA), and OsO_4_. Then, the epoxy resin (Epon)-embedded cell was serially sectioned at 50 nm-thickness, and the sections were imaged by a field-emission scanning EM (shown as EM imaging). The serial electron micrographs were three-dimensionally (3D) reconstructed for correlation with LM images. **(B)** The 3D reconstruction of mitochondria and MERCS from electron micrographs were merged with a live-fluorescence image (LM image). The ER within 15 nm of mitochondria was defined as MERCS. The boxed regions in the right upper panel are shown at higher magnification in the corresponding lower panels. **(C)** Representative images from Live-3D CLEM. The z projections of mitochondria (white line) and MERCS (yellow) in EM were overlaid with fluorescence images (Tomm20-iRFP; cyan, MERCdRED; magenta). **(D)** Comparison of MERCdRED signals and EM-identified MERCS. Proximity-mapped images of MERCdRED and MERCS were generated from (C). Each pixel value represents the average intensity of a 60-pixel (300 nm) square area centered on that pixel. The resulting MERCS images were further processed using Gaussian blurring. Chunk-averaged images of MERCdRED and MERCS were generated by averaging the proximity-mapped images within each 50-pixel (250 nm) square chunk. The signal intensities of each MERCdRED and MERCS were normalized to arbitrary units (AU). **(E)** The accuracy and precision of MERCdRED signals in colocalizing with EM-identified MERCS were assessed from the chunk-averaged images in (D). The chunks with an intensity above 0.15 were classified as positive, and the numbers of true positives, true negatives, false positives and false negatives were calculated in the MERCdRED chunk-averaged images (Original) and chunk-randomized images (Randomized) corresponding to the MERCS chunk-averaged images. The number of chunks, *#1*; n = 107, *#2*; n = 57, *#3*; n = 34, *#4*; n = 55.

### The ER-mitochondria tether PDZD8-FKBP8 protein complex is required for MERCdRED signals on mitochondria

To investigate whether MERCdRED fluorescence decreases upon inhibiting MERCS formation, we conducted the depletion of the MERCS-tethering complex in MERCdRED cells. It has been demonstrated that PDZD8 directly interacts with FKBP8, a protein localized to the outer mitochondrial membrane, and this protein complex functions as a critical tether for MERCS formation (Hirabayashi et al., 2017; Nakamura et al., 2025) (**Fig. 3A**). In the MERCdRED cells, the PDZD8 gene was conditionally and completely deleted 42 hours after treatment with 4-hydroxy (4-OH) tamoxifen (**Fig. 3A, B**). In addition, the FKBP8 gene was knocked down by infection of lentivirus expressing FKBP8-targeting short-hairpin RNA (shRNA). Consistent with roles of PDZD8 and FKBP8 previously revealed by EM analyses (Nakamura et al., 2025), deletion of either PDZD8 or FKBP8 significantly reduced MERCdRED signal intensity on mitochondria compared to that in control cells, while FKBP8 depletion in PDZD8-deficient cells had no additive effect (**Fig. 3C, D**). These demonstrate that the MERCdRED signal reflects the MERCS area in live cells (**Fig. 3C, D**).

**Figure 3.**
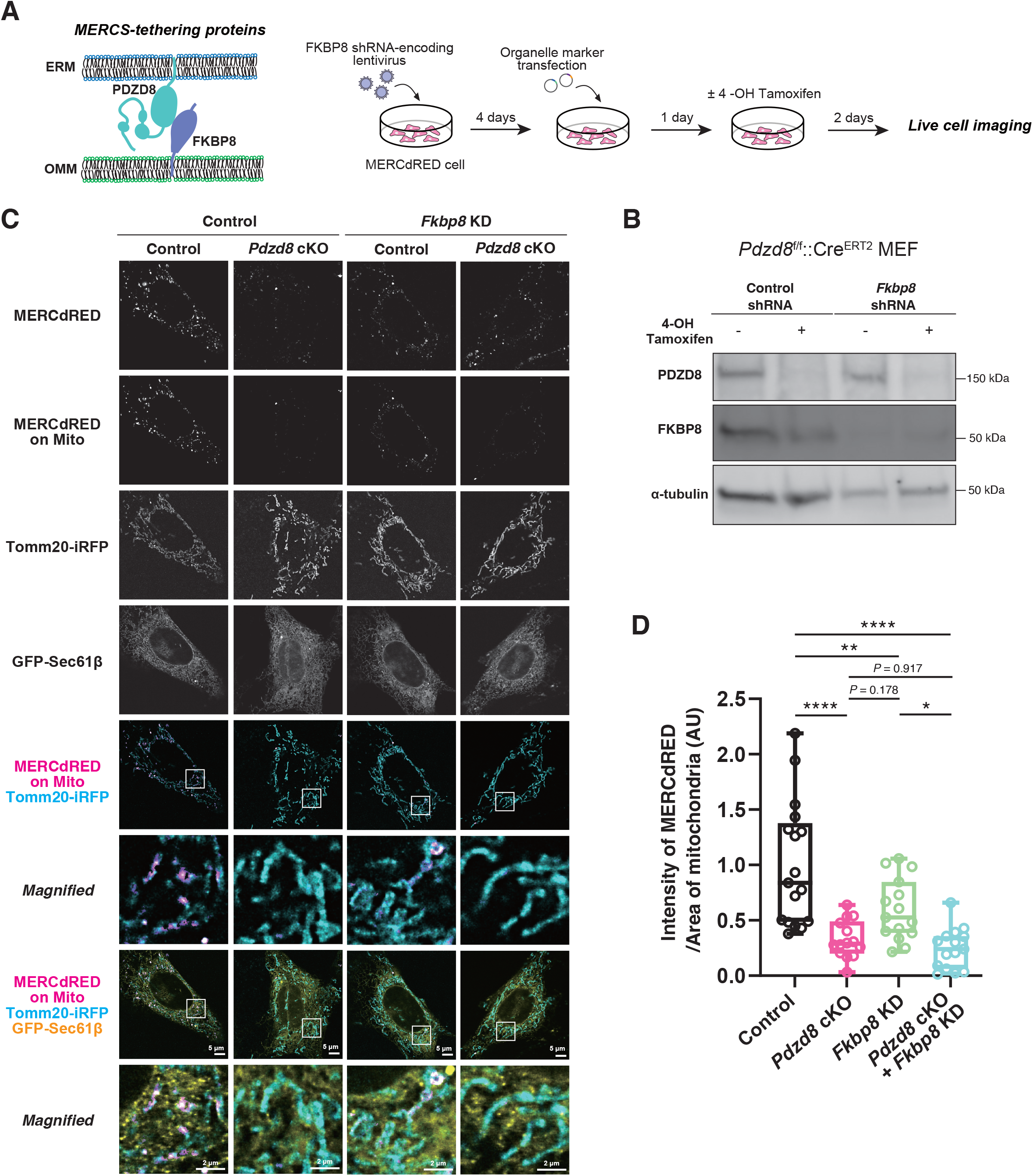
Deletion of PDZD8-FKBP8 tethering complex significantly reduced MERCdRED fluorescence. **(A)** Schematic representation depicting the depletion of the MERCS-tethering protein complex PDZD8 and FKBP8 in MERCdRED cells. *Fkbp8* gene was knocked down using lentivirus-mediated shRNA infection, and *Pdzd8* gene was knocked out by treating cells with 1 μM 4-hydroxy (4-OH) tamoxifen, which induces Cre/loxP-dependent conditional knockout. **(B)** Immunoblot analysis of *Pdzd8*^f/f^::Cre^ERT2^ MEFs infected with lentivirus carrying control shRNA or *Fkbp8* shRNA, treated with or without 0.5 μM 4-OHT. Cell lysates were subjected to immunoblotting with antibodies to PDZD8, FKBP8, and α-tubulin. **(C)** Representative live images of MERCdRED cells with or without depletion of *Pdzd8* and/or *Fkbp8*. GFP-Sec61β (yellow) or Tomm20-iRFP (cyan) was used as an ER marker or a mitochondrial marker, respectively. *Pdzd8* conditional knockout was indicated as *Pdzd8* cKO, and *Fkbp8* knockdown was shown as *Fkbp8* KD. The boxed regions of the top panels are shown at higher magnification in the corresponding lower panels. **(D)** Quantification of the MERCdRED intensity on mitochondria in images obtained as described (C). The data are presented as individual points on box plots, with the center indicating the median, and the 25th and 75th percentiles represented by the box. Whiskers extend to the minimum and maximum values. n = 17, 14, 15, 15 cells for control + control, *Pdzd8* cKO + control, control + *Fkbp8* KD, and *Pdzd8* cKO + *Fkbp8* KD, respectively, from two independent experiments. Statistical analysis was performed using Tukey’s multiple comparisons test. *\*\*\*\*P* < 0.0001, *\*\*P* < 0.01, *\*P* < 0.05.

### MERCdRED signals reveal MERCS dynamic properties in living cells

Widely employed approaches for investigating MERCS, such as EM analysis, isolation of mitochondria-associated membranes, and proximity ligation assay (PLA) require cell fixation (Scorrano et al., 2019). Therefore, by leveraging the ability to visualize MERCS in living MERCdRED cells, we investigated their dynamics by live confocal imaging. A total of 17 frames were acquired at 0.5 Hz in 20 cells, and MERCdRED-positive puncta on mitochondria were tracked across these images (**Fig. 4A, B, Supplementary movie 1**). More than 90% of MERCdRED-positive puncta had areas smaller than 0.1 μm^2^, with the median of all puncta being 0.0321 μm^2^, consistent with the MERCS areas previously defined by EM analysis and single-particle tracking of the ER-localized tethering protein VAPB (Obara et al., 2024). While the speed of individual MERCdRED puncta showed considerable variation, those with larger areas (greater than 0.05 μm^2^) moved significantly more slowly than ones with smaller areas (less than 0.05 μm^2^) (**Fig. 4C, D**). Importantly, newly emerging MERCdRED signals, corresponding to newly formed MERCS, were also observed (**Fig. 4A**, arrowheads). These findings, the dynamic formation and size-dependent motility of MERCS, demonstrate that live imaging in MERCdRED cells provides an effective platform for investigating MERCS dynamics.

**Figure 4.**
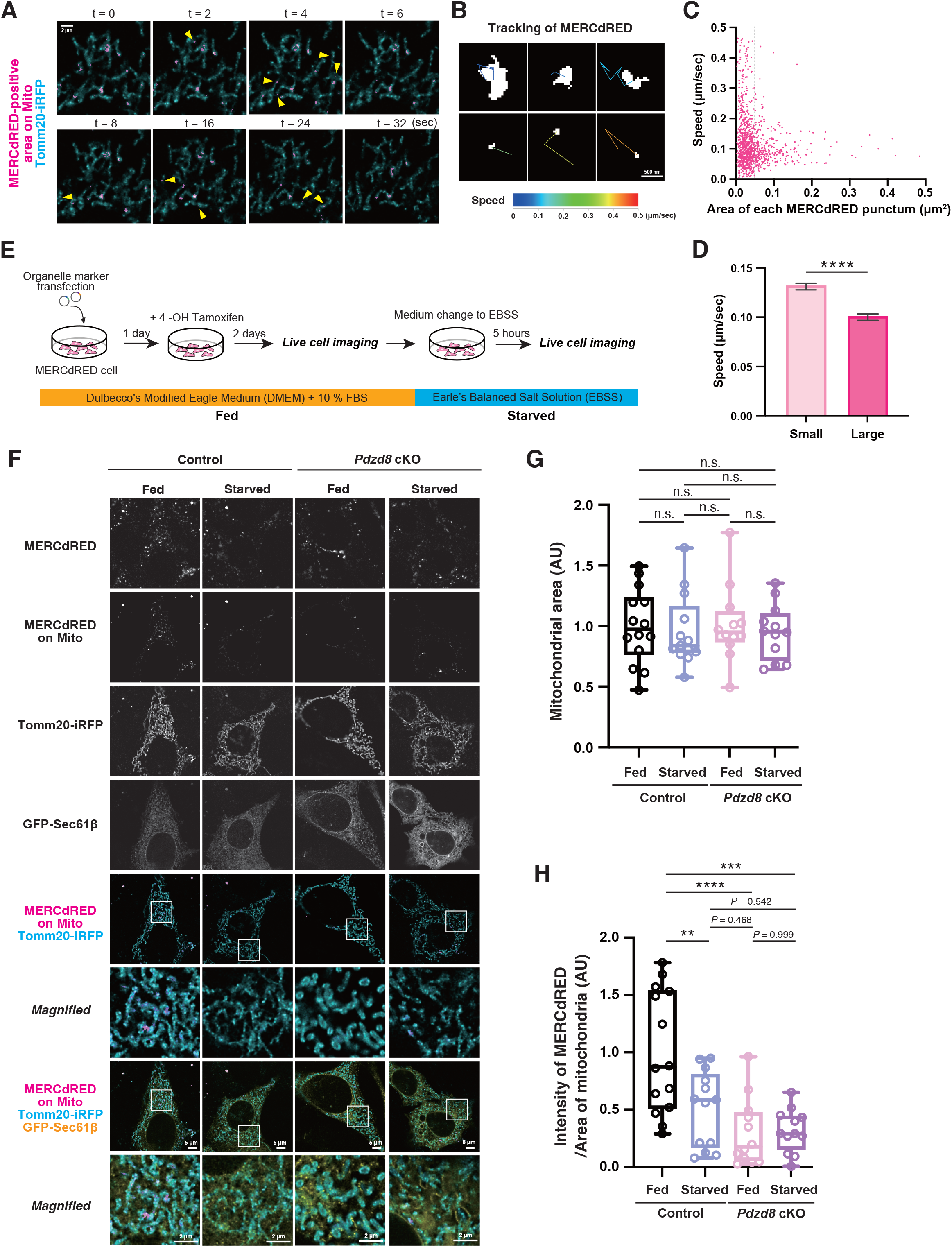
Nutrient starvation reduced PDZD8-mediated MERCS formation. **(A)** Representative images from time-lapse imaging shown as Supplementary movie 1. Images were obtained at 0.5 Hz for 32 seconds (17 frames). MERCdRED-positive areas on mitochondria were shown in magenta, and Tomm20-iRFP (cyan) was used as a mitochondrial marker. Yellow arrowheads indicate newly emerging MERCdRED signals that appeared in the subsequent time frame relative to the previous representative frame. Data are representative of 20 cells from three independent experiments. **(B)** Representative tracking images of MERCdRED-positive puncta shown in Supplementary movie 1. Tracking images were generated using TrackMate (ImageJ plugin) across the images obtained in (A). The colors of trajectories indicate the speeds of MERCdRED-positive puncta (shown in white). **(C)** Relationship between the area and the speed of each MERCdRED punctum. The average speed and area of each punctum over the entire observation period were calculated from tracking images obtained in (B). **(D)** The average speed of small puncta (area < 0.05 μm^2^; left side of the dotted line in (C)) and large puncta (area > 0.05 μm^2^; right side of the dotted line in (C)) was calculated. Data are means ± s.e.m. of 762 small puncta and 300 large puncta in 20 cells from three independent experiments. Statistical analysis was performed using the two-tailed Mann-Whitney test. **(E)** Schematic representation depicting the nutritional starvation in the MERCdRED cells. The cells were maintained in DMEM supplemented with 10% FBS and 1% penicillin-streptomycin in fed condition, whereas they were incubated in EBSS for 5 hours in starved condition. **(F)** Representative images from live-cell imaging of MERCdRED cells in the fed or starved condition. GFP-Sec61β (yellow) or Tomm20-iRFP (cyan) was used as an ER marker or a mitochondrial marker, respectively. The boxed regions of the top panels are shown at higher magnification in the corresponding lower panels. **(G)** Quantification of the mitochondrial area in images obtained as described (F). The data are presented as individual points on box plots, with the center indicating the median, and the 25th and 75th percentiles represented by the box. Whiskers extend to the minimum and maximum values. n = 14, 13, 12, 12 cells for fed + control, starved + control, fed + *Pdzd8* cKO, and starved + *Pdzd8* cKO, respectively, from two independent experiments. Statistical analysis was performed using Tukey’s multiple comparisons test. n.s., not significant. **(H)** Quantification of the MERCdRED intensity on mitochondria in images obtained as described (F). The data are presented as individual points on box plots, with the center indicating the median, and the 25th and 75th percentiles represented by the box. Whiskers extend to the minimum and maximum values. n = 14, 13, 12, 12 cells for fed + control, starved + control, fed + *Pdzd8* cKO, and starved + *Pdzd8* cKO, respectively, from two independent experiments. Statistical analysis was performed using Tukey’s multiple comparisons test. *\*\*\*\*P* < 0.0001, *\*\*\*P* < 0.001, *\*\*P* < 0.01.

### Nutritional changes modulate MERCS formation in a PDZD8-dependent manner

Finally, we investigated how MERCS dynamically responds to nutritional changes using the MERCdRED cell. The cells were maintained in a nutrient-rich medium (Dulbecco’s modified Eagle medium (DMEM) supplemented with 10% fetal bovine serum (FBS)) and subjected to starvation by culturing them in an amino acid- and serum-free, glucose-containing medium (Earle’s Balanced Salt Solution (EBSS)) for five hours (**Fig. 4E**). Strikingly, while the ratio of mitochondrial area in cells was not significantly changed (**Fig. 4F, G**), MERCdRED signals on mitochondria were significantly reduced upon starvation (**Fig. 4F, H**). In contrast, when cells were treated with 4-OHT to deplete PDZD8, PDZD8-deficient cells showed no significant changes in MERCdRED signals under the same starvation conditions (**Fig. 4F, H**). Consequently, these results suggest that PDZD8 plays a crucial role in the nutrient-dependent MERCS formation.

Although several fluorescence- or chemiluminescence-based dimerization-dependent systems have been used for MERCS visualization in living cells (Abrisch et al., 2020; Cieri et al., 2018; Csordás et al., 2010; Harmon et al., 2017; Hertlein et al., 2020; Naon et al., 2016; Naón et al., 2023; Sakai et al., 2021; Shiiba et al., 2025; Tashiro et al., 2020; Vallese et al., 2020; Yang et al., 2018), some of them face difficulties in tracking reversible MERCS formation and providing reliable quantification. In this study, the combination of the reversible association of ddFPs, stable gene expression, and single-cell cloning enabled live cell tracking of MERCS formation and quantification of MERCS areas with relatively small variance. In addition, unlike most previously reported systems, we have examined the extent of the membrane gap MERCdRED system detects using the CLEM and validated that MERCdRED can visualize MERCS with a 15 nm gap between membranes. The ability to track individual MERCdRED signals enables a detailed analysis of turnover, size, and movement of MERCS, thereby providing a deeper understanding of their dynamic properties. This system could be applied to observe MERCS changes in various cell types and cellular contexts, not only in response to nutritional changes.

Nutrient starvation has been shown to promote mitochondrial elongation in both cultured cells and mouse liver (Gomes, Benedetto, et al., 2011; Gomes, Di Benedetto, et al., 2011; Parlakgül et al., 2024; Rambold et al., 2011), likely as an adaptive response to balance energy demand and fuel supply (Liesa & Shirihai, 2013; Wai & Langer, 2016). While MERCS have been known as a site of regulating mitochondria structure through fission and fusion (Abrisch et al., 2020; Friedman et al., 2011; Ji et al., 2017), conflicting findings have been reported regarding the impact of nutrient supply changes on MERCS formation in the liver in obese mouse models and in cell lines (Arruda et al., 2014; Cieri et al., 2018; Parlakgül et al., 2024; Sakai et al., 2021; Shiiba et al., 2025; Theurey et al., 2016; Tubbs et al., 2014; Yang et al., 2018). Our results in mouse fibroblast cells support the hypothesis that MERCS formation is enhanced under nutrient-rich conditions. Since this contradicts previous studies reporting the opposite effect using other fluorescence-based or chemiluminescence-based systems, further investigation is required to clarify the influence of different MERCS detection systems, as well as differences in culture conditions and cell types. Furthermore, we revealed that nutritional enrichment promotes MERCS formation via the PDZD8-FKBP8 complex, which is responsible for regulating mitochondrial complexity (Nakamura et al., 2025). Taken together, our data suggest that nutritional starvation reduces the PDZD8-FKBP8 complex-dependent MERCS formation, leading to mitochondrial elongation. It is plausible that AMP-activated protein kinase (AMPK), which is activated in response to glucose depletion and facilitates glutaminolysis via PDZD8 (Li et al., 2024), along with other nutrient-sensing pathways, may contribute to the regulation of MERCS. Further studies are required to elucidate the precise molecular mechanisms underlying this process.

## Materials & Methods

### Cell culture

*Pdzd8*^f/f^::Cre^ERT2^ MEFs (Nakamura et al., 2025) and 293T cells (BRC) were maintained with Dulbecco’s Modified Eagle Medium (DMEM; FUJIFILM Wako Pure Chemical Corporation, catalog no. 043-30085) supplemented with 10% fetal bovine serum (FBS; MP Biomedicals, catalog no.2917346), and 1% penicillin-streptomycin (Gibco, catalog no. 15140-122) at 37°C under 5% CO_2_. For live cell imaging, phenol red-free DMEM (FUJIFILM Wako Pure Chemical Corporation, catalog no. 040-30095) supplemented with 4 M L-Glutamine (Gibco, catalog no. 25030149), 1 M Sodium Pyruvate (Gibco, catalog no. 11360070), 10% FBS and 1% Penicillin-Streptomycin was used. For starvation, cells were cultured in Earle’s Balanced Salt Solution (EBSS; Merck, catalog no. E3024) at 37°C under 5% CO_2_ for 5 hours.

### Plasmids

For FUW-Tomm20-ggsg x4-GB-P2A-RA-ggsg x3-Sec61β, the Tomm20-GB sequence was amplified by PCR from pTOM20-ddGFP-B (Addgene, catalog no. #40291) with the pair of primers 5’-agc tca agc ttc gaa ttc atg gcc tcg aga atg gtg gg -3’ and 5’-gag gag tga att gcg gcc gct tac ttg tac cgc tcg tcc a -3, and subcloned into the EcoRI and NotI sites of pCIG vector (kindly provided from Dr. Franck Polleux) with In-Fusion HD cloning kit (Takara Bio), resulting in pCAG-Tomm20-GB. The RA-Sec61β sequence was amplified by PCR from RA-NES (Addgene, catalog no. #61019) and pAc-GFP-Sec61β (Addgene, catalog no. #62008) with the pair of primers 5’-cga gct caa gct tcg aat tca tgg tga gca aga gcg agg a -3’ and 5’-tct gag gct agc ctt gta cag ctc gtc cat gc -3’, 5’-caa ggc tag cct cag atc tat gcc tgg tcc -3’ and 5’-gga gtg aat tgc ggc cgc cta cga acg agt gta ctt gcc c -3’, respectively, and subcloned into the EcoRI and NotI sites of pCIG vector with In-Fusion HD cloning kit, resulting in pCAG-RA-Sec61β. After that, the sequence of GGSG x 3 linker (ggt ggt tct ggt ggt ggt tct ggt ggt ggt tct ggt aag) was inserted prior to NheI sites in pCAG-RA-Sec61β, resulting in pCAG-RA-ggsg x3-Sec61β. Then, the Tomm20-GB-P2A sequence and the RA-ggsg x3-Sec61β sequence were amplified by PCR from pCAG-Tomm20-GB or pCAG-RA-ggsg x3-Sec61β, with the pair of primers 5’-ggg atc cac cgg tcg gta ccg cct cct ccg agg aca aca -3’ and 5’-gtc tcc agc ctg ctt cag cag gct gaa gtt agt agc tcc gct tcc ctt gta ccg ctc gtc cat g -3’ or 5’-gct aag cag gct gga gac gtg gag gag aac cct gga cct atg gtg agc aag agc gag ga -3’ and 5’-gga gtg aat tgc ggc cgc cta cga acg agt gta ctt gcc c -3’, respectively, and subcloned into the KpnI and NotI sites of pCAG-Tomm20-GB, resulting in pCAG-Tomm20-GB-P2A-RA-ggsg x3-Sec61β. The Tomm20-GGSG x 4 sequence was amplified by PCR from pCAG-Tomm20-GB-P2A-RA-ggsg x3-Sec61β with the pair of primers 5’-gcc gaa ttc atg gcc tcg aga atg gtg ggt cg -3’ and 5’-gcc acc ggt cca cca ctc cct cca cct gaa cct cca cct gag cct ccc ccg cta ccg cct tcc aca tc -3’, and subcloned into the EcoRI and AgeI sites of pCAG-Tomm20-GB-P2A-RA-ggsg x3-Sec61β with In-Fusion HD cloning kit, resulting in pCAG-Tomm20-ggsg x4-GB-P2A-RA-ggsg x3-Sec61β. Finally, the Tomm20-ggsg x4-GB-P2A-RA-ggsg x3-Sec61β sequence was amplified by PCR from pCAG-Tomm20-ggsg x4-GB-P2A-RA-ggsg x3-Sec61β with the pair of primers 5’-gcc gaa ttc atg gcc tcg aga atg gtg gg -3’ and 5’-tcg act tac ttg tac act acg aac gag tgt act tgc cc -3’, and subcloned into the EcoRI and BsrGI sites of FUW vector (Nakamura et al., 2025) with Ligation high Ver.2 (Toyobo), resulting in FUW-Tomm20-ggsg x4-GB-P2A-RA-ggsg x3-Sec61β.

For pCAG-Tomm20-iRFP, the iRFP sequence was cut out from pN1-G3BP1-iRFP (kindly provided from Dr. Yukiko Gotoh) and subcloned into the AgeI and NotI sites of pCAG-Tomm20-GB with Ligation High ver.2 (Takara). pLKO-shFKBP8 and pLKO-scramble were prepared as described previously (Nakamura et al., 2025).

### Lentivirus production

293T cells were co-transfected with shuttle vectors (FUW-Tomm20-ggsg×4-GB -P2A-RA-ggsg×3-Sec61b), HIV-1 packaging vectors Delta8.9 and VSV-G envelope glycoproteins, or shuttle vectors (pLKO-shFKBP8 or pLKO-scramble), LP1, LP2, and VSV-G using FuGENE transfection reagent (Promega, catalog no. E2311). At 24 hours post transfection, the media were exchanged with 8 mL of fresh DMEM supplemented with 10% FBS, and 1% penicillin-streptomycin, and 24 hours later, supernatants were harvested, spun at 500 g to remove debris and filtered through a 0.45 μm filter (Sartorius). The filtered supernatant was concentrated to 125 μL using an Amicon Ultra-15 (molecular weight cut-off 100 kDa) centrifugal filter device (Merck Millipore), which was centrifuged at 4,000 g for 60 minutes at 4°C. Then, 100 μL of viral supernatants was added to each 6-well dish containing MEFs.

### Establishment of MERCdRED cell line by single cell cloning using limited dilution assay

*Pdzd8*^f/f^::Cre^ERT2^ MEFs were infected with lentivirus encoding FUW-Tomm20-ggsg×4-GB-P2A-RA-ggsg×3-Sec61b. After 24 hours of incubation, the cells were washed twice with PBS and then transferred to fresh DMEM. The cells were expanded for one week and subsequently plated into a 96-well plate at a density of 0.5 cells per well. After 10 days, several proliferating clones were selected, and red fluorescence signals were assessed by live-cell imaging. Among these, the cell line exhibiting the highest red signal was selected as “MERCdRED cells” for subsequent experiments.

### Live-cell imaging of MERCdRED cells

MERCdRED cells were transfected with plasmids encoding Tomm20-iRFP and GFP-Sec61β as a mitochondrial marker or an ER marker respectively, by Lipofectamine LTX reagent with Plus reagent (Thermo Fisher). For FKBP8 knockdown, the cells were infected with lentivirus encoding FKBP8-targeting shRNA or control shRNA 4 days prior to plasmid transfection. After 4 hours incubation with plasmids, the cells were washed twice with PBS and then plated into 35 mm glass bottom dishes (Cellvis) in phenol red-free full DMEM. For PDZD8 conditional knockout, 1 μM 4-hydroxy (4-OH) tamoxifen was supplemented in medium. The cells were then imaged after 42 hours incubation. During the imaging, cells were maintained at 37°C under 5% CO_2_ in an incubation chamber (Okolab). Images were acquired on a Nikon Ti2 Eclipse microscope with an A1R confocal, a CFI Plan Apochromat Lambda D 100X Oil (NA 1.45), a laser unit (Nikon; LU-N4, 405, 488, 561, and 640 nm), and filters (525/50nm, 595/50nm, 700/75 nm for 488 nm, 561 nm, and 640 nm respectively). All equipment was controlled via NIS-elements software. Optical sectioning was performed at Nyquist for the longest wavelength. The resulting images were deconvoluted with NIS-elements (Nikon).

### Correlative light-electron microscopy

MERCdRED cells transfected with plasmids encoding Tomm20-iRFP and EGFP-Sec61β were plated in gridded coverslip-bottom dishes (custom made, based on IWAKI 3922-035; coverslips were attached inverted side), precoated with carbon by a vacuum coater and then coated with poly-d-lysine (Merck, catalog no. P0899). After live imaging, the cells were fixed with 2% paraformaldehyde and 0.5% glutaraldehyde (Electron Microscopy Sciences) in 0.1 M phosphate buffer (pH 7.4) at 37 °C for 1 hours and washed with 0.1 M phosphate buffer. The cells were then replaced in 2.5% glutaraldehyde in 0.1 M phosphate buffer for 4 days at 4 °C. After washing with 0.1 M phosphate buffer, the cells were post-fixed with 1% OsO_4_ (Electron Microscopy Sciences), 1.5% potassium ferrocyanide (Fujifilm Wako pure chemical corporation, catalog no. 161-03742) in a 0.05 M phosphate buffer for 30 minutes. After being rinsed 3 times with H_2_O, the cells were stained with 1% thiocarbohydrazide (Sigma-Aldrich) for 5 minutes. After being rinsed with H_2_O for three times, the cells were stained with 1% OsO_4_ in H_2_O for 30 minutes. After being rinsed with H_2_O for two times at room temperature and three times with H_2_O at 50°C, the cells were treated with Walton’s lead aspartate (0.635% lead nitrate (Sigma-Aldrich), 0.4% aspartic acid (pH 5.2, Sigma-Aldrich)) at 50°C for 20 minutes. The cells dehydrated with an ascending series of ethanol (15 minutes each in 50% on ice, 70% on ice, and 10 minutes each in 90%, 95% ethanol/H2O at room temperature, 10 minutes in 100% ethanol 4 times at room temperature) were embedded in epoxy resin (Epok812) by covering the gridded glass with a resin-filled beam capsule. Epok812 resin was made by mixing 7.5 g of MNA (Oken), 13.7 g of Epok812 (Oken), 3.8 g of DDSA (Oken), and 0.2 g of DMP-30 (Oken). Polymerization was carried out at 42 °C for 12 hours and 60°C for 72 hours. After polymerization, the gridded coverslip was removed, and the resin block was trimmed to a square of approximately 150 to 250 μm. The block was sectioned using an ultramicrotome (EM UC7, Leica) equipped with a diamond knife (Ultra JUMBO 45 degree, DiATOME) to cut 50 nm thick sections. The serial ultra-thin sections were collected on the cleaned silicon wafer strip and imaged with a scanning electron microscope (JSM-IT800SHL; JEOL). Imaging was done at 1 kV accelerating voltage, 39 pA beam current, 2,560 × 1,920 frame size, 6.2 mm working distance, 12.8 × 9.6 μm field of view (pixel size is 5 nm) and 1.41 μs dwell time, using the Scintillator Backscattered Electron Detector.

### Image analysis

The images taken by confocal microscopy were processed with ImageJ (NIH). For quantification of MERCdRED signals, all analyses were conducted using custom python-based programs. The mitochondrial area was defined as a region of interest (ROI) using binarizing signals of Tomm20 created by OpenCV’s threshold function, and then the sum of MERCdRED intensity at ROIs divided by the area of ROIs was calculated. Images with values exceeding ± 2 standard deviations from the mean were excluded as outliers in each experiment. For CLEM analysis, the electron micrographs were stitched by Stitch Sequence of Grids of Images Plugin and aligned using the Linear Stack Alignment with Scale Invariant Feature Transform (SIFT) plugin, implemented in ImageJ. Mitochondria and ERs in the vicinity of mitochondria in electron micrographs were semi-automatically annotated using PHILOW software (Suga et al., 2023). ER regions within 3 pixels (= 15 nm) of the mitochondrial periphery were defined as MERCS. Reconstructing segmented images of the electron micrographs to 3D images and overlaying it with fluorescence images were conducted using Imaris software (Bitplane). The z projection of electron micrographs was created using ImageJ. Proximity-mapped images and chunk-averaged images were created using custom python-based programs. In proximity-mapped images, each pixel value represents the average intensity of a 60-pixel (300 nm) square area centered on that pixel, and the resulting MERCS images were further processed using gaussian blurring. Chunk-averaged images were generated by averaging the proximity-mapped images within each 50-pixel (250 nm) square chunk. The randomization of the MERCdRED chunk list was conducted using Excel (Microsoft). For time-lapse imaging of MERCdRED, data were analyzed by the TrackMate plugin, implemented in ImageJ, with the mask detector and the simple LAP tracker (Linking max distance; 1.0 μm, Gap-closing max distance; 1.0 μm, Gap-closing max frame gap;1). For Fig. 4C, D, MERCdRED puncta with tracking durations of 10 seconds or less (i.e., 5 frames or fewer) that spanned either the initial time point (t = 0 sec) or the final time point (t = 32 sec) were excluded from the analysis. All statistical analyses were performed in Prism 10 (GraphPad Software).

### Immunoblotting

Cells were lysed with a solution containing 20 mM Hepes-NaOH (pH 7.5), 150 mM NaCl, 0.25 M sucrose, 1 mM EDTA, 0.1% SDS, 0.5% Sodium deoxycholate, 0.5 % NP-40, 1 mM Na_3_VO_4_, cOmplete Mini Protease Inhibitor Cocktail (Roche) and Benzonase (25 U/ml). Insoluble pellets and supernatants were separated by centrifugation at 15,000 × g at 4 °C for 15 minutes. The supernatants were boiled with 1 °ø Laemmli’s sample buffer containing 10% mercaptoethanol at 98 °C for 5 minutes. The cell lysates were fractionated by SDS-PAGE on a 4-15 % gradient gel (Bio-rad) and the separated proteins were transferred to a polyvinylidene difluoride membrane (Merck). The membrane was incubated first with primary antibodies for 24 hours at 4 °C and then with HRP–conjugated secondary antibodies (GE Healthcare) for 1 hour at room temperature. After a wash with TBS-T (50 mM Tris-HCl (pH 8.0), 150 mM NaCl, and 0.05 % Tween 20), the membrane was processed for detection of peroxidase activity with chemiluminescence reagents (100 mM Tris-HCl (pH 8.5), 1.25 mM Luminol, 0.2 mM P-Coumaric Acid, 0.01 % H_2_O_2_) and the signals were detected by Image Quant LAS4000 instrument (GE Healthcare). Primary antibodies; anti-PDZD8 (Hirabayashi et al., 2017; 1:500), anti-FKBP8 (R and D systems, MAB3580; 1:500), anti-α-tubulin (Sigma-Aldrich, T6188; 1:1,000).

## Acknowledgments

We thank members of the Hirabayashi lab for constructive discussions.

## Author Contributions

S.A.-I. and Y.H. conceptualized the project. Y.H. supervised the project. S.A.-I., K.N., T.N., and Y.H. designed and performed the experiments. S.A.-I. and Y.H. wrote the paper with the help of the rest of the authors.

## Competing interests

The authors declare no competing interests.

## Funding

This work was supported by JSPS KAKENHI under Grant Number JP20H04898 (Y.H.), JP22H05532 (Y.H.), JP24H01269 (Y.H.), JP21J00490 (S. A-I.), AMED under Grant number JP21wm0525015 (Y.H.), the Naito Foundation (S. A-I.),

---

**Supplementary movie 1 Time-lapse imaging of MERCdRED signals**

Time-lapse imaging was conducted at 0.5 Hz for 32 seconds (17 frames) in MERCdRED cells. MERCdRED-positive areas on mitochondria were shown in magenta, and a mitochondrial marker Tomm20-iRFP was shown in cyan. Tracking images were generated using TrackMate (ImageJ plugin). The colors of trajectories represent the speeds of MERCdRED-positive puncta (shown in white), as indicated in Figure 4B.

## Notes

### Competing Interest Statement

The authors have declared no competing interest.

